# Widespread imprinting of transposable elements and young genes in the maize endosperm

**DOI:** 10.1101/2020.04.08.032573

**Authors:** Sarah N Anderson, Peng Zhou, Kaitlin Higgins, Yaniv Brandvain, Nathan M Springer

## Abstract

Fertilization and seed development is a critical time in the plant life cycle, and coordinated development of the embryo and endosperm are required to produce a viable seed. In the endosperm, some genes show imprinted expression where transcripts are derived primarily from one parental genome. Imprinted gene expression has been observed across many flowering plant species, though only a small proportion of genes are imprinted. Understanding the rate of turnover for gain or loss of imprinted expression has been complicated by the reliance on single nucleotide polymorphisms between alleles to enable testing for imprinting. Here, we develop a method to use whole genome assemblies of multiple genotypes to assess for imprinting of both shared and variable portions of the genome using data from reciprocal crosses. This reveals widespread maternal expression of genes and transposable elements with presence-absence variation within maize and across species. Most maternally expressed features are expressed primarily in the endosperm, suggesting that maternal de-repression in the central cell facilitates expression. Furthermore, maternally expressed TEs are enriched for maternal expression of the nearest gene. Read alignments over maternal TE-gene pairs indicate fused transcripts, suggesting that variable TEs contribute imprinted expression of nearby genes.

## Main Text

Imprinted genes showing parent-of-origin based patterns of expression were first identified in maize^1^ and have since been identified in a variety of flowering plants. In plants, imprinted expression is primarily observed in the endosperm, which is a nutritive tissue of the seed that is formed when the diploid central cell is fertilized by one of the two sperm cells delivered by the pollen tube. The central cell is epigenetically distinct from most vegetative cells in the plant due to DNA demethylation targeted primarily to Transposable Elements (TEs) ^7–9^. This demethylation acts as a primary imprint that distinguishes the female and the male alleles in the endosperm. Maternal and paternal alleles are further distinguished through differential accumulation of histone modifications such as H3K27me3 ^10,11^ which often marks the maternal allele of paternally expressed genes (PEGs) while maternally expressed genes (MEGs) often show differences in DNA methylation alone ^12^.

Imprinting has been studied at the genomic level in many plant species ^2–6^. While some genes with conserved imprinting across species contribute to the establishment of imprinting ^13^, several studies have observed substantial turnover of imprinting for many genes, either within a single species or across species ^14,15^. However, understanding the rate of turnover and the source of the imprinted expression pattern has been challenging due in part to methodological inconsistencies across studies and the limitations of available SNPs for allele calls. In Arabidopsis, applying consistent methods and cutoffs across studies reduces apparent variability in imprinting calls ^16,17^, however many genes cannot be assessed due to a lack of informative SNPs. A lack of SNPs can be due to identical sequence or unalignable regions resulting from large structural changes or presence-absence variation (PAV) of whole genes or features. In maize, many genes and TEs exhibit PAV among genotypes ^18–20^. This limits the ability to use SNP-based allele-specific expression analyses to study imprinting, especially for transposons and variable genes. In this study, we develop an alternative approach that relies upon comparisons of expression in reciprocal crosses to assess the imprinting of both conserved and variable genes and TEs across maize genotypes with whole genome assemblies, revealing imprinting for many transposable elements and variable genic sequences.

Reciprocal crosses for every pairwise contrast between three maize genotypes with whole genome assemblies (B73 ^21^, W22 ^22^, and PH207 ^23^) were performed, and 14 days after pollination, endosperm was isolated in triplicate for RNA-sequencing (Table S1). Two approaches were applied to identify imprinted expression (Figure 1A). The traditional approach for calling imprinting uses Single Nucleotide Polymorphisms to call Allele Specific Expression (SNP-ASE) followed by comparison of biases across reciprocal crosses (methods). The SNP- ASE ratio is calculated by assigning SNP-containing reads to one allele and determining the proportion of informative reads from each allele, providing an estimate of the expression of two alleles within a single sample. We developed and implemented an alternative approach where reads are aligned to concatenated genome files and the Reciprocal Expression Ratio (RER) was calculated to describe the ratio of expression for features in each genome when inherited maternally versus paternally. Unlike SNP-ASE, the RER is a comparison of expression of a feature in reciprocal crosses and cannot be calculated for a single sample. Calculations of RER rely on the ∼15% of reads that map uniquely to a single location in the concatenated genomes (Table S1). While many reads map equally well to both genomes and are therefore discarded, unique mapping reads are only found in places of the genome with variants distinguishing the alleles (SNPs or indels) or in regions unique to one genome. After assigning unique reads to features including genes and TEs using HTseq, RER was calculated by dividing the expression level (RPM) when inherited maternally by the sum of expression when maternally or paternally inherited. Given that endosperm is composed of two copies of the maternal genome and one copy of the paternal genome, the null expectation for a transcript’s expression is that it will be twice as highly expressed when inherited from the maternal parent compared to the paternal parent. For both SNP-ASE and RER, the average value representing a biparentally expressed gene is 0.67, allowing direct comparison of the methods. A comparison of SNP-ASE and RER reveals general agreement between these two approaches for genes that could be analyzed with SNPs, with the majority of genes expressed at the ratio expected by dosage (Figure 1B). Many of the genes showing disagreement between methods in Figure 1B result from genotype-biased expression which exhibits a strong bias in SNP-ASE for a single sample but doesn’t result in bias for RER (Figure S1). To further assess accuracy of RER, expression patterns for three MEGs and three PEGs with conserved imprinting status in maize, rice, and Arabidopsis ^4^ were assessed (Figure 1C, Table S2). In most cases with informative reads, clear parental bias in the expected direction was observed for all genes (Figure 1C).

**Figure 1:**
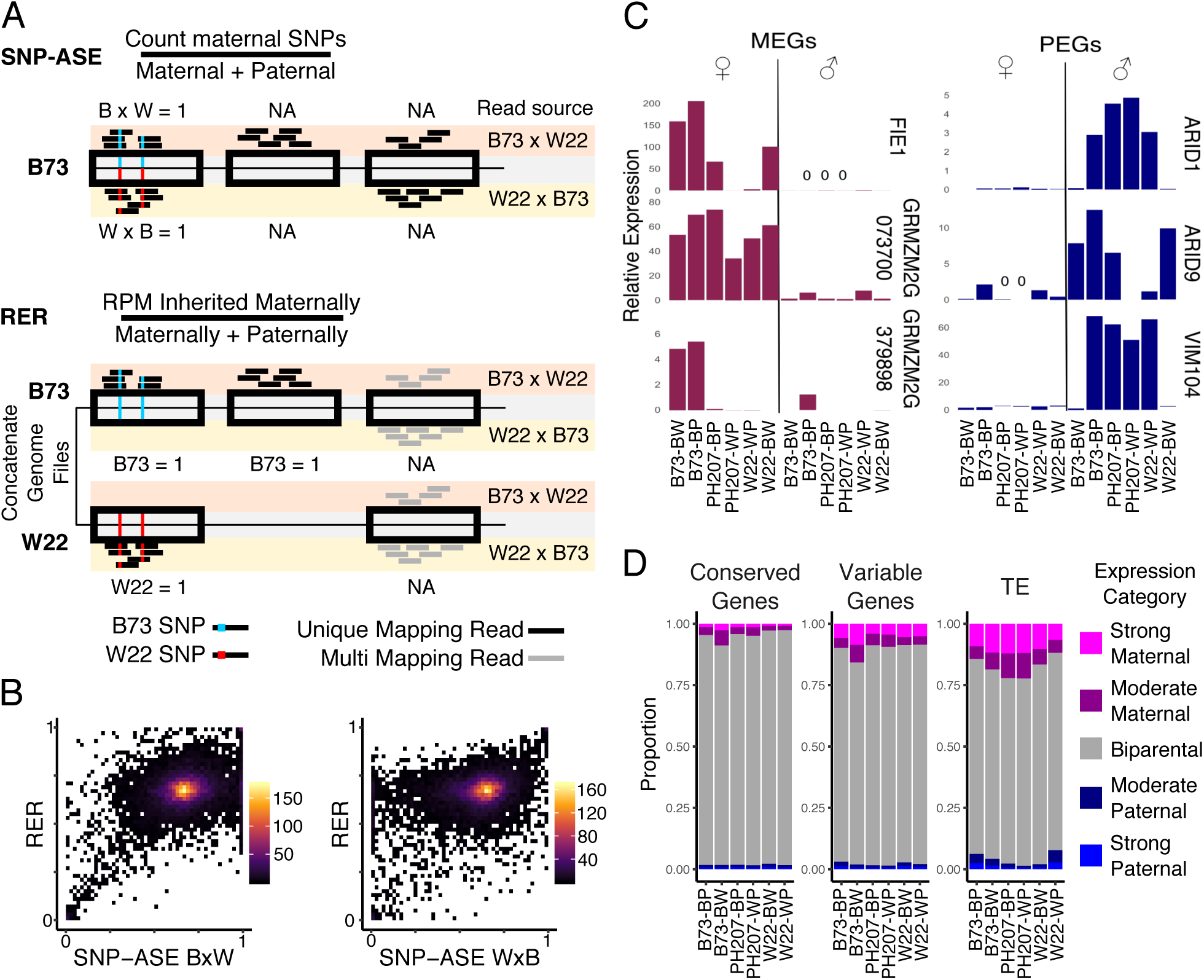
Assessing imprinted expression pattern in maize. A) The method for defining imprinting using SNP- ASE versus RER. in SNP-ASE, reads mapping to a SNP-corrected reference genome are assigned to alleles based on the SNP supported. In RER, reads are assigned to a concatenated reference genome and retained at unique positions. Both methods can be used to assess imprinting for shared genes distinguished by SNPs, but only RER can assess imprinting for PAV features. B) Comparison of SNP-ASE and RER for B73 genes accessible using both methods in the B73 x W22 cross, with values plotted showing the average across three biological replicates. SNP-ASE is assessed for each direction of reciprocal crosses separately while RER is calculated with reciprocals. The heat represents the number of genes in each pixel of the plot. C) Expression across all contrasts for genes with conserved imprinting in maize, rice, and Arabidopsis (Waters et al 2011) using the RER method. Bar height represents the mean expression across replicates. Symbols above the plot show whether the gene was inherited maternally or paternally. Gene IDs for these genes are listed in Table S2. D) The distribution of RER values for different features across contrasts. RER cutoffs for strong maternal and strong paternal are > 0.9 and < 0.1, respectively, and cutoffs for moderate maternal and paternal are > 0.8 or < 0.2, respectively.

While both methods can be used to define imprinting for shared genes distinguishable by SNPs, only the RER method can capture imprinting for portions of the genome that exhibit PAV. This provides new opportunities to study parent-of-origin biased gene expression for TEs and variable genes. The distribution of RER values was assessed across contrasts for different feature types (Figure S2), and the proportion of each set that showed parentally-biased expression was summarized based on RER (Figure 1D). This revealed that across all contrasts, genes conserved within maize rarely exhibit parent-of-origin biased expression (Figure 1D, Figure S2). On average, < 3% of expressed genes that are present in all three maize genotypes in this study show a strong parental bias (Figure 1D). For genes that are variable among maize lines, a higher proportion (> 6%) of expressed genes show high parental bias, with this set representing genes that are accessible using RER but not SNP-ASE. Strikingly, > 11% of expressed TEs show a strong parental bias, with the majority of strongly biased TEs expressed maternally (Figure 1D).

In order to identify imprinted transcripts, we applied the lfcThreshold option within DESeq2 to test for significance (adjusted p-value < 0.05) over the expected 2:1 gene dosage across reciprocals using three biological replicates. To increase the stringency of imprinting calls, significant hits were further filtered by RER values. Maternally Expressed Genes (MEGs) and Maternally Expressed TEs (matTEs) were filtered for RER > 0.9, while Paternally Expressed Genes (PEGs) were filtered for RER < 0.1. It can be difficult to remove all maternal tissues when isolating endosperm tissue and therefore it is important to limit potential false-positive calls of maternal expression that may result from genes expressed in the maternal seed coat ^24^. Previously published RNA-seq data ^25^ was used to filter out genes whose maternal expression could result from seed coat contamination rather than maternal expression in the endosperm. Pericarp-preferred genes were defined where the mean expression in pericarp was >2-fold higher than the expression in endosperm (Figure S3). After implementing these criteria and filters, we identified an average of 182 total imprinted genes across all hybrid combinations, with an average of 112 MEGs and 70 PEGs in each (Figure 2A).

**Figure 2:**
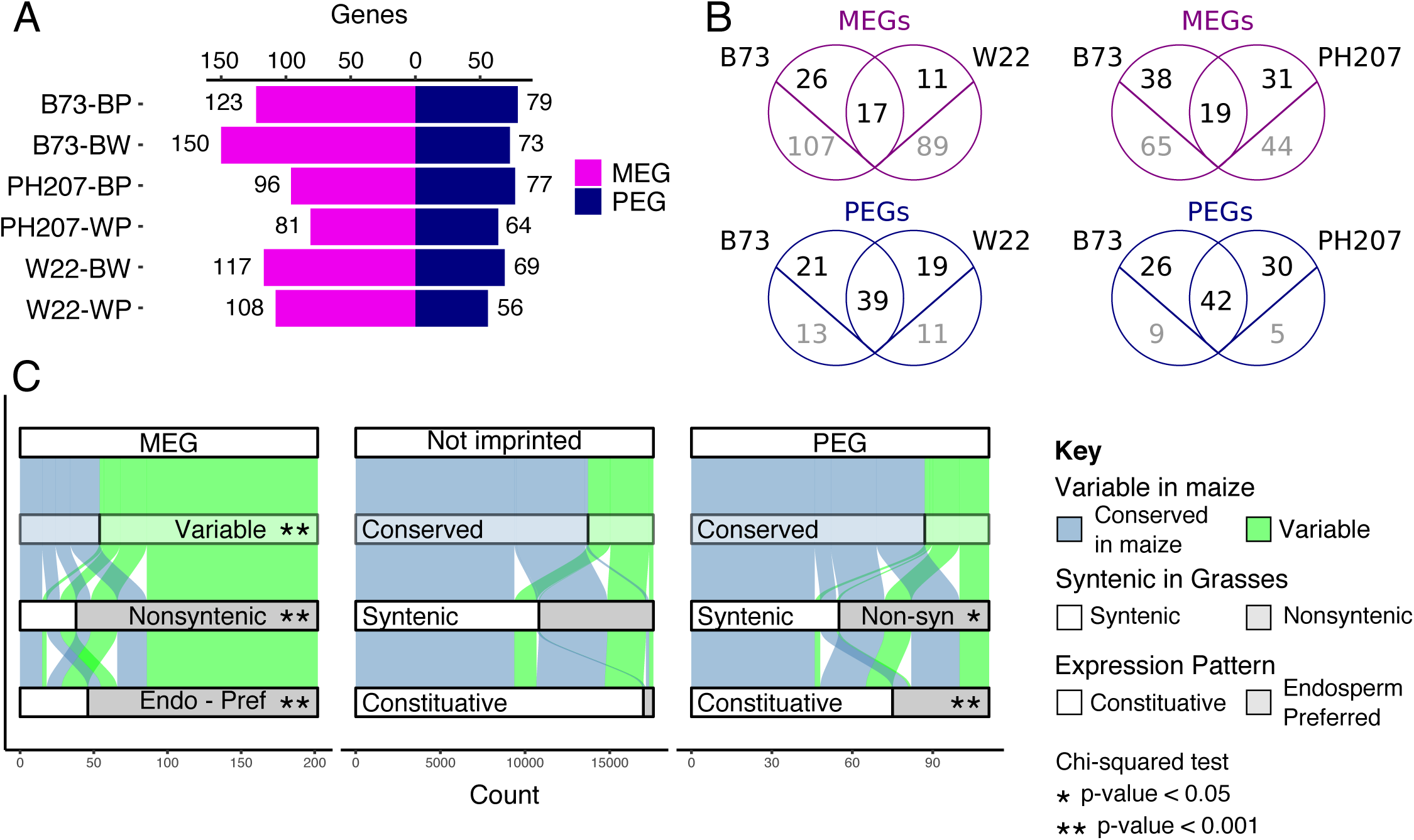
Imprinting of genes defined by RER. A) The number of imprinted genes identified across contrasts using the RER method (see methods). MEGs are shown in magenta and PEGs are shown in blue. B) The overlap between imprinted genes across pairwise contrasts. Genes that are shared between genotypes that could be assessed for imprinting are shown in black above the line while imprinted genes unique to one genome are shown in gray below the line. C) Comparison of features for MEGs, PEGs, and non-imprinted B73 genes. Genes are defined as conserved when they are shared with all genotypes present in this study, syntenic when a syntenic ortholog exists in sorghum, rice, foxtail millet, and brachypodium, and endosperm-preferred if expression is primarily restricted to the endosperm (Figure S4). Asterisks denote significance relative to the Not Imprinted set (chi-squared test).

The imprinted genes discovered in each genome were compared to assess the consistency of imprinting. A comparison of imprinted features in the B73 x W22 reciprocal hybrid endosperm tissue identifies 17 MEGs, 39 PEGs, and 4 matTEs that were consistently imprinted in both genomes (Figure 2B, 3B). A subset of the genes that do not exhibit consistent imprinting are shared between the two genomes. For example, there are 26 MEGs observed only in B73 and 11 only observed in W22 despite the fact that both genomes retain a syntenic ortholog for these genes. For the majority of these shared genes with variable imprinting, the lack of overlap is due to cutoff stringency or lack of coverage rather than true turnover of imprinting (Figure S4). There are many additional cases where imprinted genes are only present in one genome. For PEGs, variable genes represent the minority of non-conserved imprinted genes, with only 13 of 34 B73 PEGs that are not imprinted in W22 variable across genomes. In contrast, for the majority of MEGs with inconsistent imprinting (i.e. 107 of 133 B73 genes in the B73 by W22 contrast), the genes themselves are absent from the other genome. Similar patterns are observed for the B73 by PH207 contrast, though a higher proportion of genes are shared in this contrast. The large number of maternally expressed transcripts with variability in maize suggests that imprinting of non-conserved elements may be far more prevalent than previously detected due to the limitations of SNP-based allele calls.

To understand additional features of imprinted genes, we focused on the B73 genes that were called imprinted in at least one contrast, which included 202 MEGs and 111 PEGs. B73 was selected as the central genotype because it has substantially more expression datasets, syntenic gene information, and functional gene annotations than other genomes. For the genes identified as imprinted, we compared several characteristics relative to genes that were expressed but were not classified as imprinted. First, genes were assessed for variability across maize inbred lines by defining conserved genes as those with syntenic orthologs in B73, W22, and PH207 and variable genes as those without a corresponding gene in at least one genome ^26^. This revealed a clear enrichment for variable genes among MEGs (p-value < 0.001, chisq test), but not PEGs, compared to genes that are not imprinted but have enough unique reads to be assessed for imprinting (Figure 2C). We then expanded our evolutionary distance and assessed how many genes in each set are syntenic with other grasses as defined by having a syntenic ortholog in sorghum, rice, foxtail millet, and brachypodium. For genes without imprinting, the majority (62%) are syntenic with other grasses. However, MEGs are highly depleted for syntenic genes (19%) and PEGs show a minor depletion (50%, p-value < 0.05, chisq test). Next, the expression pattern across B73 development was assessed using published RNA-seq data ^25^. Since imprinting can arise from either silencing of one parental allele specifically in the endosperm or de-repression of one parental allele in the endosperm, the pattern of expression across tissues was defined as either constitutive or endosperm-preferred (see methods, figure S4). While only 3% of non-imprinted genes are expressed preferentially in the endosperm, 77% of MEGs and 32% of PEGs show this expression pattern (Figure 2C, p-value < 0.001, chisq test). Many of the MEGs (38%) have no assigned GO term, a 2.8-fold enrichment compared to genes that are not imprinted (p-value < 0.001, chisq test). Since TEs are a common source of new genes and a driver of gene content variation among maize lines, we intersected our imprinted genes with annotated TEs, identifying 26 MEGs and 1 PEG completely within an annotated transposable element. While MEGs and PEGs are annotated as genes in the B73v4 annotation, transcription of a locus does not imply the creation of a functional gene product. While evolutionarily conserved genes with synteny to other grasses may be the best candidates for real genes capable of conferring phenotypes ^27^, variable genes can be important for functions such as disease resistance ^28^.

To further investigate the imprinting of TEs themselves, the RER method was used to define imprinted TEs, with an average of 95 matTEs identified across contrasts (Figure 3A). There are a small number of paternally expressed TEs, however these were excluded from further analyses due to the low number detected and potential technical complications (Figure 3A, S2). Consistent with the large amount of TE variability among genotypes, the majority of imprinted TEs were unique to one genome (Figure 3B). There are 145 maternally expressed TEs in B73 relative to at least one other genotype, including 72 LTR retrotransposons, 52 Helitrons, 9 TIR transposons, and 2 LINEs (Figure 3C). The vast majority of these TEs (93%) represent specific TE insertions that are polymorphic among the three maize genotypes ^20^. Given the high tissue-specificity of TE expression observed previously ^29^, the tissue-specific expression patterns for matTEs were also assessed. We found that 92% of matTEs are expressed preferentially in the endosperm, suggesting that imprinting is established through de-repression of the maternal allele preferentially in the endosperm and that this is the only stage of development for expression of these elements (Figure 3C, S5). Since TE families have the potential for coordinated expression responses among members, the families for matTEs were assessed. matTEs are in 84 families, with only one Helitron family containing more than 5 imprinted elements. This family, DHH00002 (DHH2), contains 44 maternally expressed members and is the only Helitron family in B73 that is predicted to have autonomous members. Since prior work has suggested that Helitrons are responsible for creating imprinting by moving PHE1 binding sites around the genome ^30^, the proportion of DHH2 Helitrons with predicted motifs was assessed (Figure S6). We found that matTEs of this family are more likely to have a binding site than elements that are not detected in our analysis, though the distribution is similar to family members that are not imprinted so it is unlikely that PHE1 sites alone are sufficient to confer imprinting of DHH2 Helitrons.

**Figure 3:**
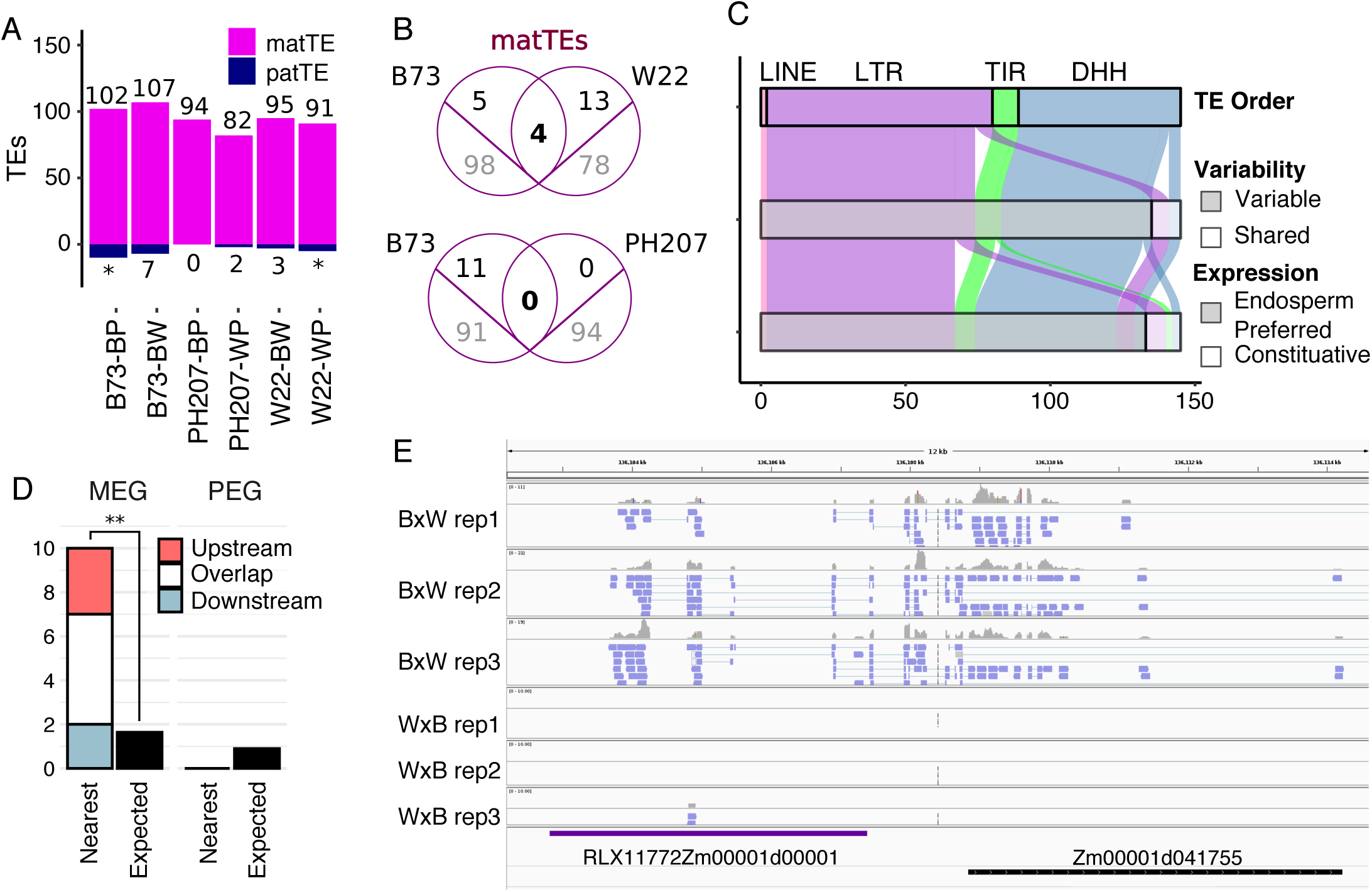
Imprinted TEs defined by RER. A) The number of imprinted TEs across contrasts. matTEs are marked in magenta and paternally expressed TEs are marked in navy. Asterisks denote contrasts where paternally expressed TEs could not be defined (Figure S2). B) The overlap between matTEs across pairwise contrasts. TEs that are shared between genotypes that could be assessed for imprinting are shown in black above the line while imprinted TEs unique to one genome are shown in gray below the line. C. Features of matTEs in B73. TE orders are abbreviated: DHH = Helitron, TIR = terminal inverted repeat transposon, LTR = LTR retrotransposon, and LINE = long interspersed nuclear element. TE variability is defined by prior work (Anderson et al. 2019). Endosperm-preferred expression is described by patterns across development (Figure S4). D) Imprinting status of closest gene to matTEs. Expected number is based on the number of MEGs and PEGs that were assessed for imprinting. ** p-value < 0.001 (binomial test) E) IGV view showing a representative example of reads aligning to a matTE near a MEG. Reads are colored by the strand of alignments, where blue = forward strand.

TEs have been proposed as a source of variation in imprinted gene expression, and this dataset allows for investigation into the relationship between imprinted genes and TEs. For every matTE in B73, the closest gene was identified and assessed for imprinting. For 13% of matTEs, the nearest gene is a MEG, which is a significant enrichment (p-value < 0.001, binomial test) and 11.6 times more common than expected based on the proportion of expressed genes that are called MEGs (Figure 3D). In contrast, there were no identified examples of matTEs where the closest gene is a PEG. There were 19 matTEs where the closest gene is one of 15 MEGs (Table S3). In the majority of cases, the TE overlapped (N = 7) or was upstream of the gene (N = 10). We identified only two cases of the TE located downstream of the gene, and one of these genes also overlapped a matTE. The asymmetry between upstream and downstream TE relationships suggests that the orientation likely matters for determining which TEs are able to influence gene expression patterning. In all cases, the developmental expression patterns of the genes and the nearby TEs match. To understand the nature of transcripts, read alignments for matTE-MEG pairs were visualized with IGV. In all cases, reads aligning to both the matTE and corresponding MEG mapped to the same strand without clear separation in read alignments, suggesting that many of these clusters may actually represent single transcripts overlapping multiple features (Figure 3E).

In summary, we developed the RER method to use information from shared and variable portions of maize whole genome assemblies to identify imprinted expression of genes and TEs in maize. This revealed imprinting of many genes that were undetectable by traditional methods that rely on diagnostic SNPs between parental alleles. The majority of maternally expressed features (genes and TEs) represent young portions of the genome that are variable within maize and non-syntenic with other grasses. We also observe strong enrichment for MEGs near maternally expressed TEs, further supporting the connection between turnover of imprinting and presence-absence variation of TEs. In mammals, imprinting in the placenta has been proposed to result from different defense mechanisms used by male and female germlines to reduce retrovirus proliferation in the germ line ^31^, and turnover of imprinting could have a similar host defense explanation in plant endosperm. In plants, there are genes with conserved imprinting across plant species that support theories of parental conflict ^32^ or dosage ^33^, however the majority of imprinted loci are variable within and across species. By studying imprinting using whole genome assemblies, we are able to better understand the turnover of imprinted expression of both shared and variable portions of plant genomes.

## Materials and Methods

### Materials

Three maize inbred lines, B73, W22, and PH207, were grown in the field in Saint Paul, MN in the summer of 2018. Reciprocal crosses between each pair of genotypes were performed. Ears were collected 14 days after pollination and endosperm was isolated using manual dissection, with approximately 10 kernels per ear pooled for each biological replicate. Paired-end, stranded RNA-seq libraries were created using the Illumina TruSeq Stranded mRNA kit and sequencing was performed with the Illumina HiSeq 2500 at the University of Minnesota Genomics Center. On average, > 45 million reads were generated per library (Table S1).

### Sequence alignments for RER

Concatenated genome files were created for each pairwise contrast of parental genomes and assemblies used included B73v4 ^21^, W22 ^22^, and PH207 ^23^. When necessary, chromosome designations were altered to ensure non-redundant sequence names across parents. Hisat2 index files were created using genome sequences only for each contrast. Gene annotations and disjoined filteredTE annotations available at https://github.com/SNAnderson/maizeTE_variation were combined by first subtracting exon regions from the TE annotations and then combining full gene and TE annotations for each genome. Concatenated annotation files were then created for each pairwise contrast using the same chromosomal designation as for the genome files. RNA-seq reads were trimmed using cutadapt ^34^ and aligned to the concatenated genomes corresponding to the parents using hisat2 ^35^. Unique-mapping reads to the concatenated genome files were then assigned to features (genes and TEs) using HTseq ^36^. Counts to each feature were normalized as reads per million using library size estimates derived from the SNP- ASE method (described below). RER for each annotation (gene and TE) was calculated by dividing the mean expression when inherited maternally by the sum of the expression when inherited maternally and paternally.

### Sequence alignments for SNP-ASE

In parallel to the above method of mapping reads, we also ran the standard, SNP-based allele specific expression pipeline by mapping reads to the B73 AGPv4 reference assembly using a variant-aware aligner HiSat2 trained with a set of known SNPs as described in ^37^. The number of reads supporting each parental genotype were used to calculate the proportion of maternal reads for each gene. For comparison across mapping methods, genes were filtered for only those with at least 10 informative reads in both methods. SNP-ASE ratios were calculated for each gene in each direction of the reciprocal cross separately by dividing the number of reads matching the maternal allele by the total number of informative reads. Genes with parent-specific expression were defined as those with a SNP maternal ratio > 0.85 in one direction and < 0.15 in the reciprocal direction.

### Defining imprinting

To define imprinted features using RER, count tables for genes and TEs in each library were loaded into R. For each of the three reciprocal crosses performed in triplicate, DESeq2 ^38^ was applied using the lfcThreshold=1 and altHypothesis=“greaterAbs” options to identify features with significant deviations from the 2:1 expected expression difference based on dosage. Each contrast includes features from both parental genomes, so maternal and paternal expression was determined by the direction of the differential expression plus the genome where the feature was annotated. Significant features were further filtered to only strong cases of imprinting where RER was > 0.9 for MEGs and matTEs and < 0.1 for PEGs. To create the final list of imprinted features, maternal features with pericarp-preferred expression were filtered out (see Tissue Dynamics).

### Tissue Dynamics

The expression profile of genes and TEs was analyzed for B73 features using previously published analysis ^29^ using data from ^25^. To filter out genes where expression is higher in the pericarp than the endosperm and could thus result in inaccurate imprinting calls ^24^, expression was compared for 14 dap seeds (the time point used in this study) and 18 dap pericarp. Genes with expression over twice as high in the pericarp over the endosperm were excluded from MEG calls. W22 and PH207 genes corresponding to genes expressed higher in the pericarp were also excluded from MEG calls. No matTEs were identified as potential contaminants using this method. Expression data across all tissues was also used to identify endosperm-preferred expression. Endosperm-preferred expression was defined as genes and TEs where the sum of expression in endosperm and wole seed libraries (26% of libraries) was more than 60% of the sum of expression across all libraries.

### Descriptors

To identify genes that are shared between genome assemblies and annotations, the file gene_model_xref_v4.txt was downloaded from MaizeGDB ^26^ on 2020/01/22. This file is B73- based and genes with a single corresponding gene in either the pairwise contrast (for venn diagrams) or in both W22 and PH207 (all other analyses) were defined as conserved in maize while remaining genes were defined as variable. This file was also used to define genes that are syntenic with other grasses, with syntenic genes being defined as any gene with a syntenic ortholog in foxtail millet, rice, brachypodium, and sorghum. To identify the nearest gene to each matTE, bedtools closest was used and distances between TE and gene were reported relative to the orientation of the gene.

## Data Availability

RNA-seq data files have been uploaded to NCBI SRA under BioProject ID PRJNA623806. Scripts and data files used to process results are available at https://github.com/SNAnderson/Imprinting2020 and https://github.com/kmhiggins/Imprinting_2020.

## Acknowledgements

We thank Peter Hermanson for technical assistance. This work was funded by grants from USDA-NIFA 2016-67013-24747 (SNA, and NMS), NSF IOS-1546899 (PZ and NMS), NSF DGE-1545453 (KH), and from the Minnesota Agricultural Experiment Station (MIN 71-068). The Minnesota Supercomputing Institute (MSI) at the University of Minnesota provided computational resources that contributed to this research.

## References

1. Kermicle, J. L. Dependence of the R-mottled aleurone phenotype in maize on mode of sexual transmission. Genetics 66, 69–85 (1970).

2. Hsieh, T.-F. et al. Regulation of imprinted gene expression in Arabidopsis endosperm. Proc. Natl. Acad. Sci. U. S. A. 108, 1755–1762 (2011).

3. Luo, M. et al. A genome-wide survey of imprinted genes in rice seeds reveals imprinting primarily occurs in the endosperm. PLoS Genet. 7, e1002125 (2011).

4. Waters, A. J. et al. Parent-of-origin effects on gene expression and DNA methylation in the maize endosperm. Plant Cell 23, 4221–4233 (2011).

5. Zhang, M. et al. Genome-wide screen of genes imprinted in sorghum endosperm, and the roles of allelic differential cytosine methylation. Plant J. 85, 424–436 (2016).

6. Hatorangan, M. R., Laenen, B., Steige, K. A., Slotte, T. & Köhler, C. Rapid Evolution of Genomic Imprinting in Two Species of the Brassicaceae. Plant Cell 28, 1815–1827 (2016).

7. Gehring, M., Bubb, K. L. & Henikoff, S. Extensive demethylation of repetitive elements during seed development underlies gene imprinting. Science 324, 1447–1451 (2009).

8. Ibarra, C. A. et al. Active DNA demethylation in plant companion cells reinforces transposon methylation in gametes. Science 337, 1360–1364 (2012).

9. Park, K. et al. DNA demethylation is initiated in the central cells of Arabidopsis and rice. Proc. Natl. Acad. Sci. U. S. A. 113, 15138–15143 (2016).

10. Weinhofer, I., Hehenberger, E., Roszak, P., Hennig, L. & Köhler, C. H3K27me3 profiling of the endosperm implies exclusion of polycomb group protein targeting by DNA methylation. PLoS Genet. 6, (2010).

11. Moreno-Romero, J., Jiang, H., Santos-González, J. & Köhler, C. Parental epigenetic asymmetry of PRC2-mediated histone modifications in the Arabidopsis endosperm. EMBO J. 35, 1298–1311 (2016).

12. Zhang, M. et al. Genome-wide high resolution parental-specific DNA and histone methylation maps uncover patterns of imprinting regulation in maize. Genome Res. 24, 167–176 (2014).

13. Luo, M., Bilodeau, P., Dennis, E. S., Peacock, W. J. & Chaudhury, A. Expression and parent-of-origin effects for FIS2, MEA, and FIE in the endosperm and embryo of developing Arabidopsis seeds. Proc. Natl. Acad. Sci. U. S. A. 97, 10637–10642 (2000).

14. Waters, A. J. et al. Comprehensive analysis of imprinted genes in maize reveals allelic variation for imprinting and limited conservation with other species. Proc. Natl. Acad. Sci. U. S. A. 110, 19639–19644 (2013).

15. Pignatta, D. et al. Natural epigenetic polymorphisms lead to intraspecific variation in Arabidopsis gene imprinting. Elife 3, e03198 (2014).

16. Wyder, S., Raissig, M. T. & Grossniklaus, U. Consistent Reanalysis of Genome-wide Imprinting Studies in Plants Using Generalized Linear Models Increases Concordance across Datasets. Sci. Rep. 9, 1320 (2019).

17. Picard, C. L. & Gehring, M. Identification and Comparison of Imprinted Genes Across Plant Species. Methods Mol. Biol. 2093, 173–201 (2020).

18. Springer, N. M. et al. Maize inbreds exhibit high levels of copy number variation (CNV) and presence/absence variation (PAV) in genome content. PLoS Genet. 5, e1000734 (2009).

19. Hirsch, C. N. et al. Insights into the maize pan-genome and pan-transcriptome. Plant Cell 26, 121–135 (2014).

20. Anderson, S. N. et al. Transposable elements contribute to dynamic genome content in maize. (2019) doi:10.1101/547398.

21. Jiao, Y. et al. Improved maize reference genome with single-molecule technologies. Nature 546, 524–527 (2017).

22. Springer, N. M. et al. The maize W22 genome provides a foundation for functional genomics and transposon biology. Nat. Genet. (2018) doi:10.1038/s41588-018-0158-0.

23. Hirsch, C. N. et al. Draft Assembly of Elite Inbred Line PH207 Provides Insights into Genomic and Transcriptome Diversity in Maize. Plant Cell 28, 2700–2714 (2016).

24. Schon, M. A. & Nodine, M. D. Widespread Contamination of Arabidopsis Embryo and Endosperm Transcriptome Data Sets. Plant Cell 29, 608–617 (2017).

25. Stelpflug, S. C. et al. An Expanded Maize Gene Expression Atlas based on RNA Sequencing and its Use to Explore Root Development. Plant Genome 9, (2016).

26. Portwood, J. L., 2nd et al. MaizeGDB 2018: the maize multi–genome genetics and genomics database. Nucleic Acids Res. 47, D1146–D1154 (2019).

27. Schnable, J. C. Genome evolution in maize: from genomes back to genes. Annu. Rev. Plant Biol. 66, 329–343 (2015).

28. Xu, X. et al. Resequencing 50 accessions of cultivated and wild rice yields markers for identifying agronomically important genes. Nat. Biotechnol. 30, 105–111 (2011).

29. Anderson, S. N. et al. Dynamic Patterns of Transcript Abundance of Transposable Element Families in Maize. G3 9, 3673–3682 (2019).

30. Batista, R. A. et al. The MADS-box transcription factor PHERES1 controls imprinting in the endosperm by binding to domesticated transposons. Elife 8, (2019).

31. Haig, D. Retroviruses and the placenta. Curr. Biol. 22, R609–13 (2012).

32. Haig, D. Coadaptation and conflict, misconception and muddle, in the evolution of genomic imprinting. Heredity 113, 96–103 (2014).

33. Dilkes, B. P. & Comai, L. A differential dosage hypothesis for parental effects in seed development. Plant Cell 16, 3174–3180 (2004).

34. Martin, M. Cutadapt removes adapter sequences from high-throughput sequencing reads. EMBnet.journal 17, 10–12 (2011).

35. Kim, D., Langmead, B. & Salzberg, S. L. HISAT: a fast spliced aligner with low memory requirements. Nat. Methods 12, 357–360 (2015).

36. Anders, S., Pyl, P. T. & Huber, W. HTSeq--a Python framework to work with high-throughput sequencing data. Bioinformatics 31, 166–169 (2015).

37. Zhou, P., Hirsch, C. N., Briggs, S. P. & Springer, N. M. Dynamic Patterns of Gene Expression Additivity and Regulatory Variation throughout Maize Development. Mol. Plant 12, 410–425 (2019).

38. Love, M. I., Huber, W. & Anders, S. Moderated estimation of fold change and dispersion for RNA-seq data with DESeq2. Genome Biol. 15, 550 (2014).

