## Supplemental Figures and Tables for "Widespread imprinting of transposable elements and young genes in the maize endosperm"

Figure S1

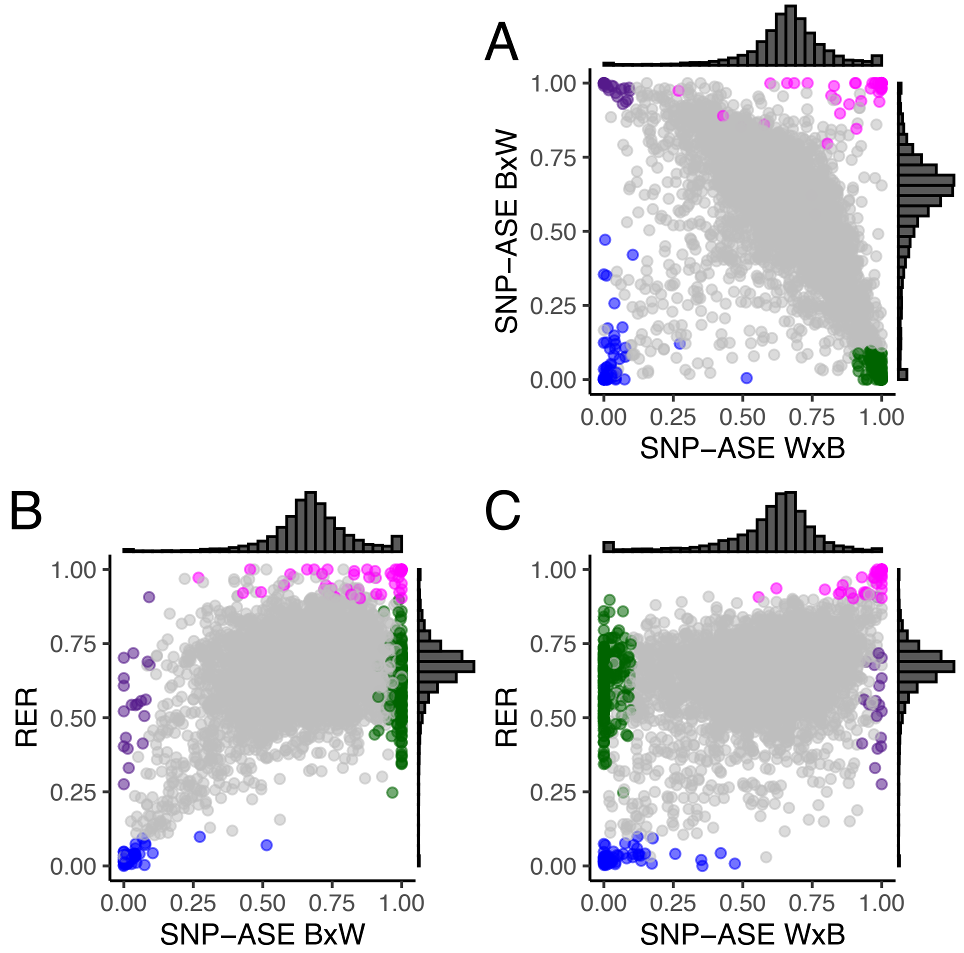


Figure S1: Comparison of SNP-ASE and RER for B73 genes accessible using both methods in the B73 x W22 cross. A) The SNP-ASE method applied across both directions of reciprocal crosses, with points colored based on imprinting calls using RER (magenta and blue) or inconsistency across reciprocals using SNP-ASE (green and purple). Colors are consistent across panels. B-C) Comparison of the two SNP-ASE values to RER. Plots are the same as shown in Figure 1B but colored to indicate imprinted genes and genes with genotype-biased expression using SNP-ASE. Genotype bias does not impact RER since the calculation is performed across reciprocals instead of across genotypes. For all panels, values plotted show the average across three biological replicates. Genes are color-coded based on expression pattern, with magenta = MEG, blue = PEG, green = B73 biased, and purple = W22 biased. Genotype bias was defined by SNP-ASE ratios > 0.85 in one direction and < 0.15 in the other direction of reciprocal crosses.

Figure S2

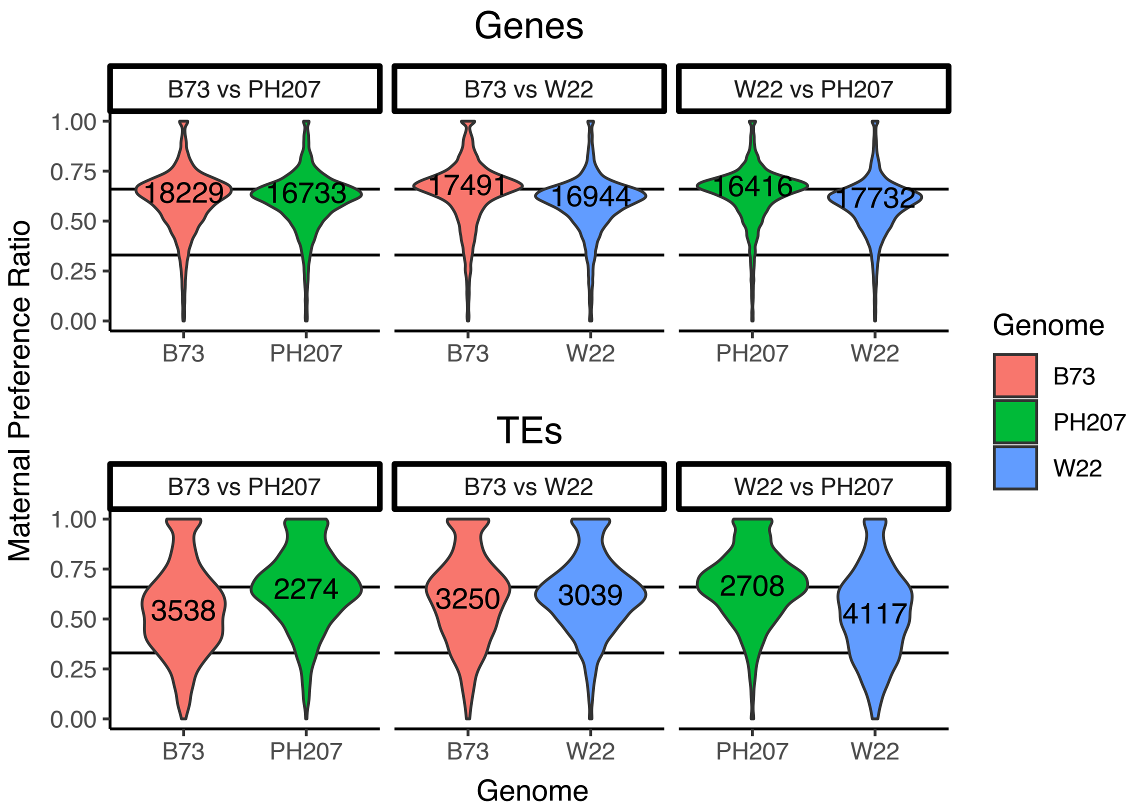


Figure S2: Distribution of RER for genes and TEs across contrasts. Most genes and TEs are expressed near the expected ratio given genome dosage (horizontal line, 0.67), with maternal and paternal expression approaching 1 or 0, respectively. TEs have a more pronounced peak of maternal expression than genes. For TEs, a bimodal peak near 0.67 and 0.33 in contrasts with PH207 suggests that some PH207 TEs may be mapping better to B73 or W22 assemblies due to poor assembly quality of PH207 in intergenic space. For this reason, paternal TEs were not counted in contrasts with PH207 in Fig 3A.

Figure S3

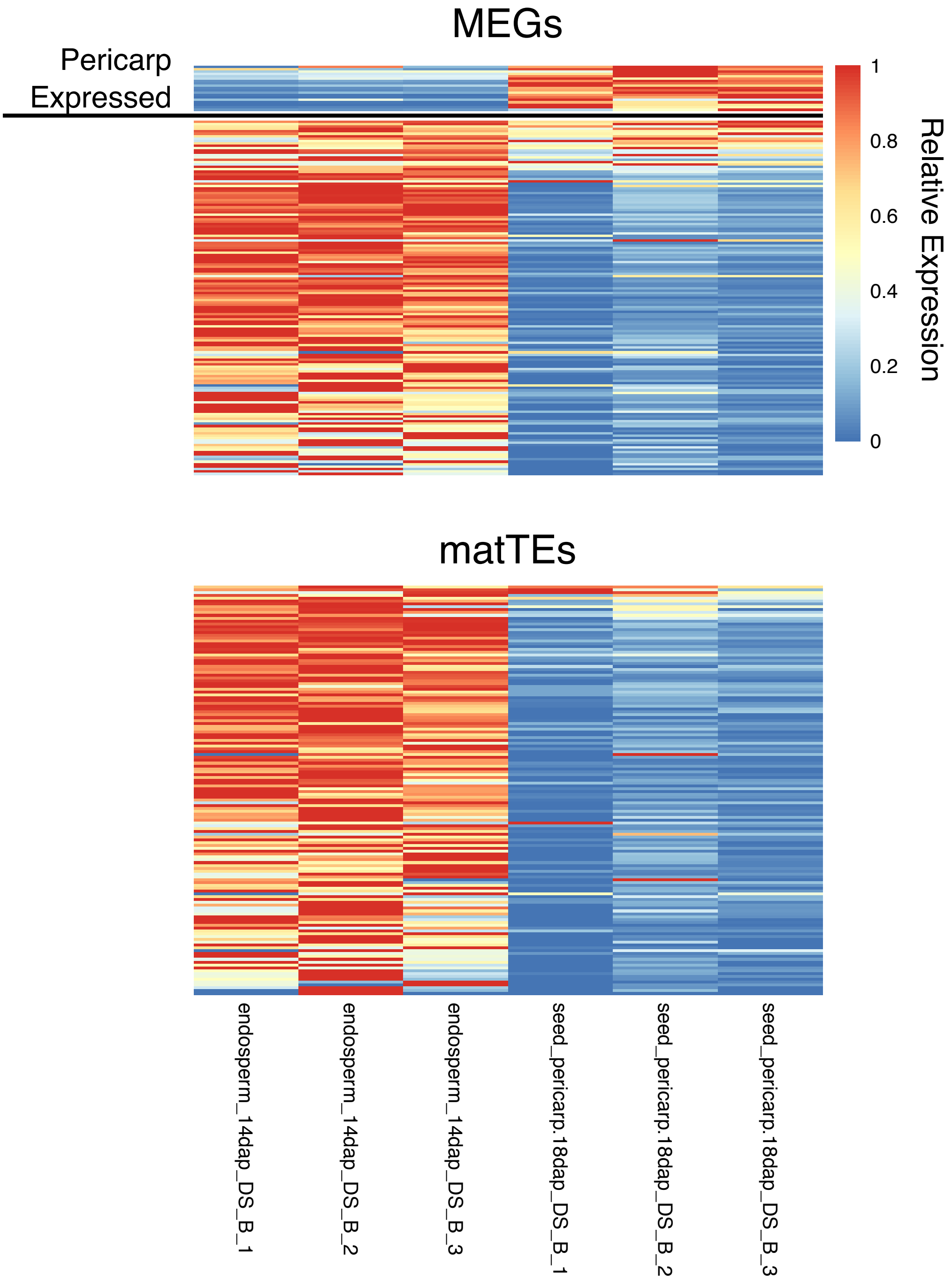


Figure S3: Expression profile of B73 MEGs and matTEs in the endosperm and pericarp using RNA-seq data from Stelpflug et al. Genes with mean expression in the pericarp > 2x the mean expression in the endosperm were filtered from the MEG list due to the potential for seed coat contamination. W22 and PH207 genes corresponding to B73 pericarp-preferred genes were also removed from MEG counts. There were no matTEs filtered out using this method. Heat of each pixel represents the expression value compared to the max in the row.

#### Figure S4

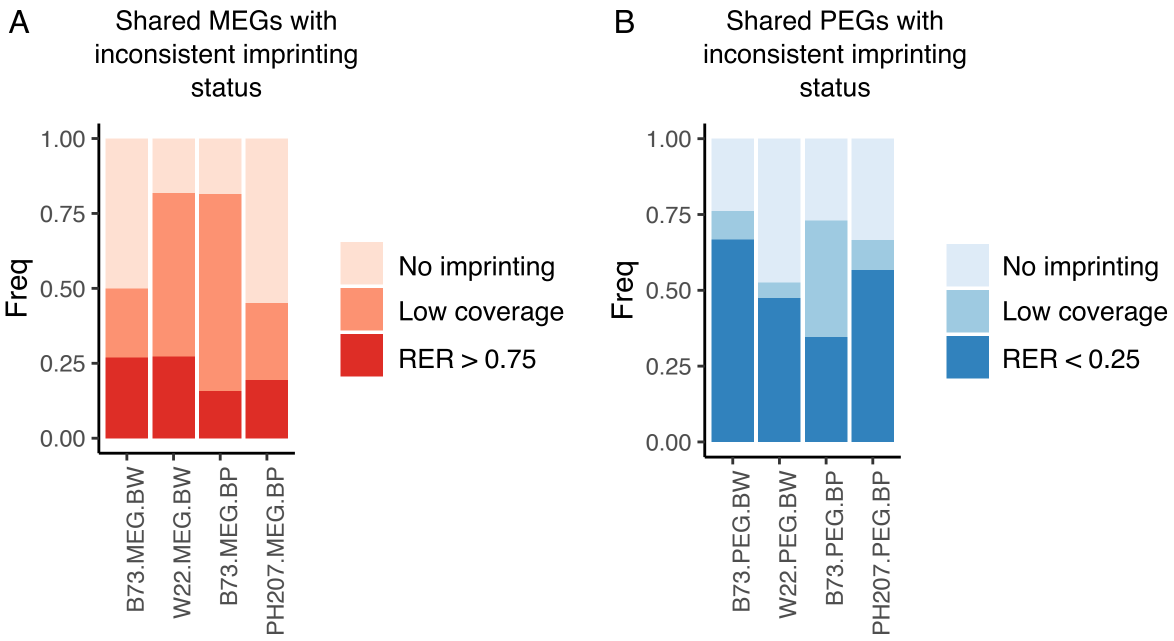


Figure S4: RER bias for genes with inconsistent imprinting in Figure 2B. For both MEGs (right) and PEGs (left), the majority of genes do not overlap due to low coverage or have RER values in the same direction as imprinted genes but failed to meet our strict statistical and/or RER threshold. X-axis labels denote the genotype where the gene is imprinted, the type of imprint, and the cross where the imprinting is variable (B = B73, W = W22, P = PH207)

Figure S5

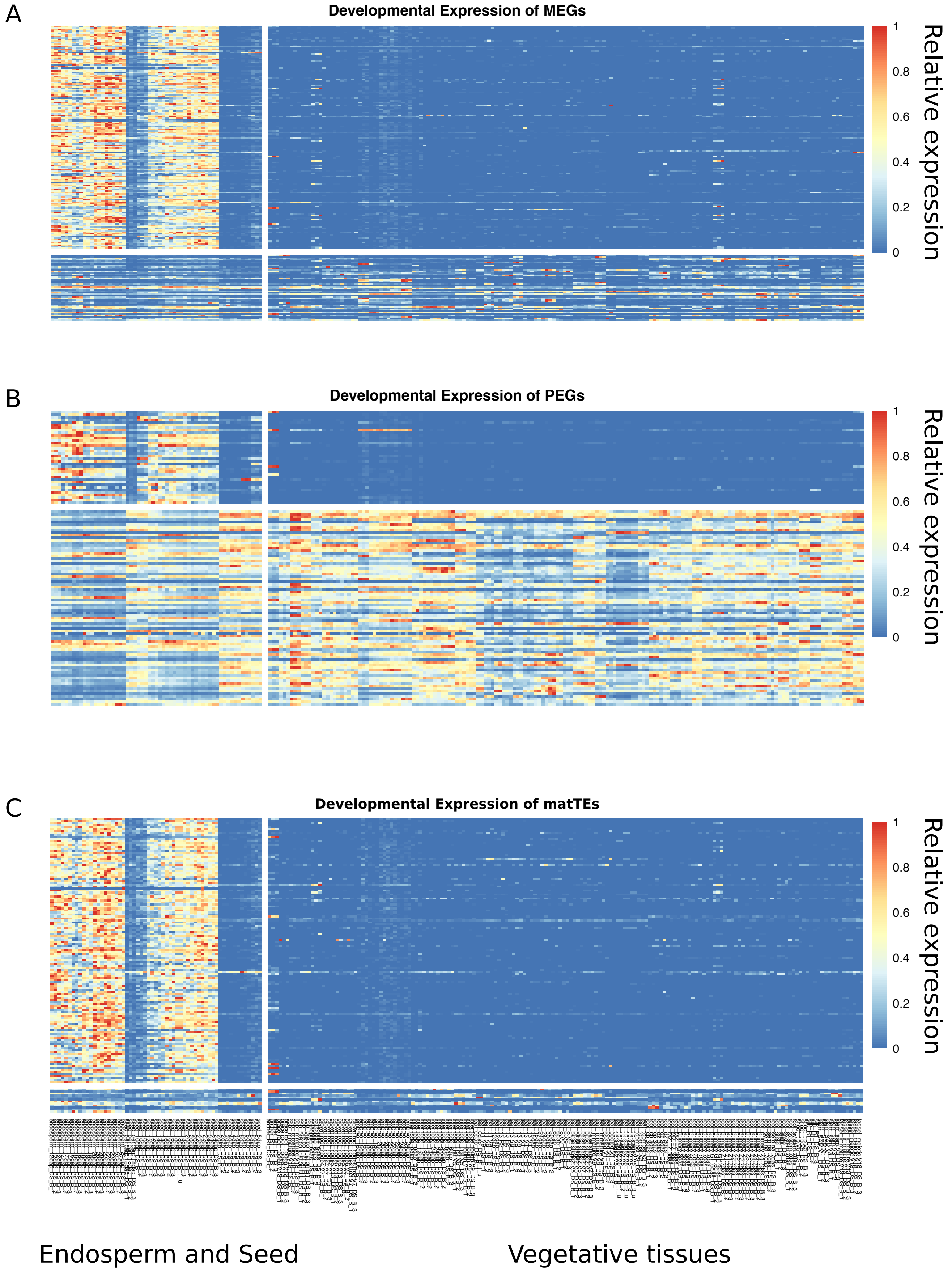


Figure S5: Expression pattern of imprinted B73 genes and TEs across development using data from Stelpflug et al. Endosperm and seed tissues are shown on the left side of the break, and all other tissues sampled are shown on the right of the break. Endosperm-preferred expression was defined where the sum of the expression across endosperm and seed libraries were more than 60% of the sum of the expression across all libraries. For each plot, endosperm-preferred features are shown above the break and constitutive features are shown below the break. Heat of each pixel represents the expression value compared to the max in the row.

Figure S6

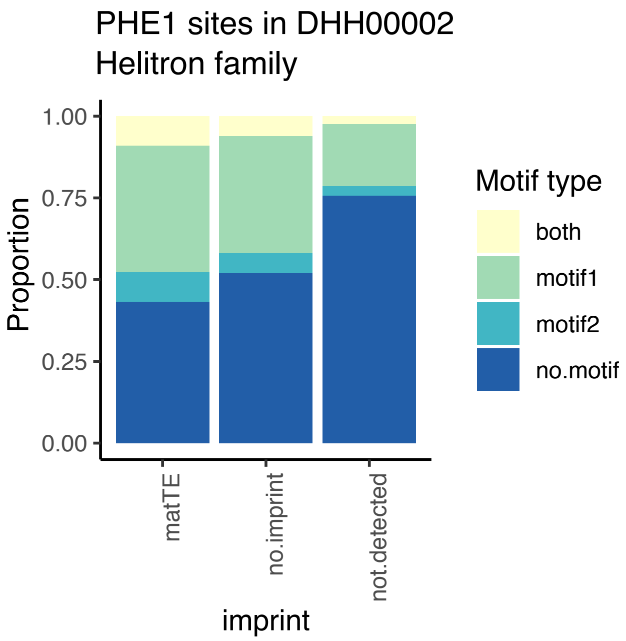


Figure S6: Presence of PHE1 binding sites within DHH2 family helitrons. Binding motifs from Batista et al. 2019 were identified in all DHH2 helitrons, and the distribution of members with zero, one, or both of the sites were determined. A similar distribution of motifs were found for matTEs and non-imprinted family members, with a lower proportion of elements with at least one motif identified in the not detected set. TEs in the not detected set include TEs that are not expressed and TEs without unique sequence that could be assessed for imprinting.

### Table S1

| Library | Reads | Percent_Unique |
|---|---|---|
| BW1 | 50912503 | 16.09983 |
| BW2 | 46955481 | 16.63385 |
| BW3 | 43297034 | 17.88011 |
| WB1 | 42341423 | 18.12524 |
| WB2 | 38350809 | 14.4479 |
| WB3 | 53936174 | 13.86724 |
| BP1 | 50287029 | 14.24699 |
| BP2 | 51260636 | 16.18052 |
| BP3 | 45525383 | 16.54365 |
| PB1 | 43670832 | 19.41905 |
| PB2 | 39706669 | 18.70737 |
| PB3 | 41024925 | 15.71401 |
| WP1 | 53203456 | 15.92903 |
| WP2 | 52838664 | 16.56664 |
| WP3 | 41023269 | 19.63248 |
| PW1 | 53900552 | 15.71085 |
| PW2 | 35267586 | 18.17527 |
| PW3 | 35654524 | 18.32236 |

Table S1: Reads and mapping statistics for RNA-seq libraries in this study

Table S2

| Gene ID | Feature | Genome | Imprinted |
|---|---|---|---|
| GRMZM2G365731 (ARID1) | Zm00001d032832 | B73 | PEG |
| AC191534.3_FG003 (VIM104) | Zm00001d019342 | B73 | PEG |
| GRMZM2G073700 | Zm00001d037209 | B73 | MEG |
| GRMZM5G866423 (ARID9) | Zm00001d032096 | B73 | PEG |
| GRMZM2G118205 (FIE1) | Zm00001d049608 | B73 | MEG |
| GRMZM2G379898 | Zm00001d027290 | B73 | MEG |
| GRMZM2G365731 (ARID1) | Zm00004b004198 | W22 | PEG |
| AC191534.3_FG003 (VIM104) | Zm00004b035225 | W22 | PEG |
| GRMZM2G073700 | Zm00004b029838 | W22 | MEG |
| GRMZM5G866423 (ARID9) | Zm00004b003587 | W22 | PEG |
| GRMZM2G118205 (FIE1) | Zm00004b020902 | W22 | MEG |
| GRMZM2G379898 | Zm00004b000042 | W22 | MEG |
| GRMZM2G365731 (ARID1) | Zm00008a004277 | PH207 | PEG |
| AC191534.3_FG003 (VIM104) | Zm00008a027204 | PH207 | PEG |
| GRMZM2G073700 | Zm00008a025023 | PH207 | MEG |
| GRMZM5G866423 (ARID9) | Zm00008a003685 | PH207 | PEG |
| GRMZM2G118205 (FIE1) | Zm00008a015599 | PH207 | MEG |
| GRMZM2G379898 | Zm00008a000045 | PH207 | MEG |

Table S2 - Gene IDs for conserved imprinted genes plotted in Figure 1C

### Table S3

| TE | gene | distance | TE.type | gene.type | order | ratio.BW.gene | ratio.BP.gene | ratio.BW.TE | ratio.BP.TE | gene.variability | gene.expression | TE.expression |
|---|---|---|---|---|---|---|---|---|---|---|---|---|
| DHH00002Zm00001d06247 | Zm00001d029042 | 0 | matTE | MEG | DHH | NA | 1 | NA | 1 | variable | endo.preferred | endo.preferred |
| RLG00003Zm00001d01354 | Zm00001d031633 | -925 | matTE | MEG | RLG | 1 | NA | 1 | 0.841323333 | variable | endo.preferred | endo.preferred |
| RLX19061Zm00001d00001 | Zm00001d023985 | 0 | matTE | MEG | RLX | 1 | 1 | 0.97095182 | 1 | variable | endo.preferred | endo.preferred |
| RLC00002Zm00001d00679 | Zm00001d026623 | -797 | matTE | MEG | RLC | 0.996352427 | 0.735195542 | 1 | 0.701963246 | conserved.maize | endo.preferred | endo.preferred |
| DHH00002Zm00001d01432 | Zm00001d004147 | 0 | matTE | MEG | DHH | 0.992924908 | 1 | 1 | 1 | variable | endo.preferred | endo.preferred |
| RLC02716Zm00001d00002 | Zm00001d005712 | -29185 | matTE | MEG | RLC | 1 | 1 | 1 | 1 | variable | endo.preferred | endo.preferred |
| RLG00003Zm00001d02516 | Zm00001d005712 | -3087 | matTE | MEG | RLG | 1 | 1 | 1 | 1 | variable | endo.preferred | endo.preferred |
| RLC00004Zm00001d02983 | Zm00001d007488 | -852 | matTE | MEG | RLC | 0.954011986 | 0.971259533 | 0.912021829 | 0.991412204 | variable | constitutive | constitutive |
| RLX11772Zm00001d00001 | Zm00001d041755 | -1453 | matTE | MEG | RLX | 1 | NA | 1 | NA | variable | endo.preferred | endo.preferred |
| DHH00002Zm00001d07691 | Zm00001d041887 | 0 | matTE | MEG | DHH | 0.982580808 | 0.846116079 | 0.986576458 | 0.961109425 | variable | endo.preferred | endo.preferred |
| RLX11447Zm00001d00001 | Zm00001d043716 | 1670 | matTE | MEG | RLX | 0.986874555 | NA | 0.945756767 | NA | conserved.maize | endo.preferred | endo.preferred |
| RLX11446Zm00001d00001 | Zm00001d043716 | 0 | matTE | MEG | RLX | 0.986874555 | NA | 0.987866795 | 0.739333538 | conserved.maize | endo.preferred | endo.preferred |
| DHH00002Zm00001d02512 | Zm00001d048646 | 0 | matTE | MEG | DHH | 1 | NA | 1 | 0.415964714 | variable | endo.preferred | endo.preferred |
| DHH00002Zm00001d02687 | Zm00001d050068 | 0 | matTE | MEG | DHH | 1 | 1 | 1 | 1 | variable | endo.preferred | endo.preferred |
| DHH00002Zm00001d08890 | Zm00001d017481 | 1078 | matTE | MEG | DHH | 0.932781433 | 1 | 1 | 0.99597214 | variable | endo.preferred | endo.preferred |
| RLG00017Zm00001dS0065 | Zm00001d038034 | -2662 | matTE | MEG | RLG | 0.914617966 | 0.776546976 | 0.956875803 | 0.817167014 | variable | constitutive | constitutive |
| RLG00001Zm00001d12334 | Zm00001d046395 | -47897 | matTE | MEG | RLG | NA | 1 | 1 | 1 | variable | endo.preferred | endo.preferred |
| RLG00003Zm00001d07748 | Zm00001d046395 | -3398 | matTE | MEG | RLG | NA | 1 | NA | 1 | variable | endo.preferred | endo.preferred |
| RLC00183Zm00001dS0162 | Zm00001d046395 | -10170 | matTE | MEG | RLC | NA | 1 | NA | 1 | variable | endo.preferred | endo.preferred |

Table S3: IDs and features of matTEs near MEGs
